## Supplementary file 1 for "Interplay between cohesin and TORC1 links chromosome segregation and gene expression to environmental changes"

**Supplementary file 1. Strains used in this study**

| strain # | genotype | Figure |
| --- | --- | --- |
| 2 | *h-* | 1A |
| 2729 | *h- mis4-G1487D* | 1A |
| 6849 | *h- mis4-G1487D pef1*Δ::*natR* | 1A |
| 12208 | *h- mis4-G1487D mip1-Q167P* | 1A |
| 11562 | *h- mis4-G1487D mip1-Y169C* | 1A |
| 10874 | *h- mis4-G1487D mip1-R401G* | 1A |
| 12226 | *h- mis4-G1487D mip1-T1007R* | 1A |
| 11558 | *h- mis4-G1487D mip1-D1018Y* | 1A |
| 11200 | *h- mis4-G1487D tor2-H2142Q* | 1A |
| 11478 | *h- psk1-13myc-hygR* | 1C |
| 12191 | *h- mip1-D1018Y psk1-13myc-kanR* | 1C |
| 12183 | *h- mip1-Y169C psk1-13myc-HygR* | 1C |
| 12252 | *h- mip1-Q167P psk1-13myc-HygR* | 1C |
| 12248 | *h- mip1-T1007R psk1-13myc-hygR* | 1C |
| 11448 | *h- mip1-R401G psk1-13myc-hygR* | 1C |
| 2 | *h-* | 1C |
| 2729 | *h- mis4-G1487D* | 1C |
| 10874 | *h- mis4-G1487D mip1-R401G* | 1C |
| 11584 | *h- mis4-G1487D mip1-Y533A-13myc-kanR* | 1C |
| 11636 | *h- mis4-G1487D mip1-13myc-kanR* | 1C |
| 2 | *h-* | 1D |
| 2729 | *h- mis4-G1487D* | 1D |
| 11002 | *h- mip1-R401G* | 1D |
| 10874 | *h- mis4-G1487D mip1-R401G* | 1D |
| 5913 | *h- eso1-H17* | 1D |
| 12318 | *h- rad21-K1* | 1D |
| 2 | *h-* | 1E |
| 2729 | *h- mis4-G1487D* | 1E |
| 10874 | *h- mis4-G1487D mip1-R401G* | 1E |
| 11420 | *h- mis4-G1487D tsc2*Δ::*kanR* | 1E |
| 12115 | *h- mis4-G1487D tsc2*Δ::*kanR mip1-R401G* | 1E |
| 2 | *h-* | 1E |
| 2729 | *h- mis4-G1487D* | 1E |
| 12175 | *h- mis4-G1487D grt1*Δ::*kanR* | 1E |
| 12179 | *h- mis4-G1487D gtr1-Q61L-kanR* | 1E |
| 12167 | *h- mis4-G1487D gtr1-S16N-kanR* | 1E |
| 12177 | *h- mis4-G1487D iml1*Δ::*kanR* | 1E |
| 12171 | *h- mis4-G1487D gcn2*Δ::*hygR* | 1E |
| 2 | *h-* | 1-sup1A |
| 2796 | *h- leu1-32 mis4-G1487D rec8-GFP-kanR* | 1-sup1A |
| 2899 | *h- leu1-32 mis4-G1487D rec8-GFP-kanR pef1-N146S* | 1-sup1A |
| 2912 | *h- leu1-32 mis4-G1487D rec8-GFP-kanR mip1-R401G* | 1-sup1A |
| 2852 | *h- leu1-32 mis4-G1487D rec8-GFP-kanR caa1-S252L* | 1-sup1A |
| 2923 | *h- leu1-32 mis4-G1487D rec8-GFP-kanR tor2-H2142Q* | 1-sup1A |
| 2 | *h-* | 1-sup1B |
| 2729 | *h-mis4-G1487D* | 1-sup1B |
| 12650 | *h- ura4-D18 mis4-G1487D tor1*Δ::*ura4+* | 1-sup1B |
| 12667 | *h- ura4-D18 tor1*Δ::*ura4+ mip1-R401G* | 1-sup1B |
| 10874 | *h- mis4-G1487D mip1-R401G* | 1-sup1B |
| 2729 | *h-mis4-G1487D* | 1-sup1B |
| 2 | *h-* | 2A-B |
| 2729 | *h- mis4-G1487D* | 2A-B-C |
| 10874 | *h- mis4-G1487D mip1-R401G* | 2A-B |
| 2729 | *h-mis4-G1487D* | 2C |
| 11326 | *h- pph3*Δ::*hygR psm3K105NK106N* | 2D |
| 11395 | *h- pph3*Δ::*kanR psm3K105NK106N mip1-R401G* | 2D |
| 12365 | *h- cdc11-GFP-kanR ndc80-GFP-kanR mis6-RFP-hygR* | 2E-G |
| 12363 | *h- cdc11-GFP-kanR ndc80-GFP-kanR mis6-RFP-hygR mis4-G1487D* | 2E-G |
| 12393 | *h- cdc11-GFP-kanR ndc80-GFP-kanR mis6-RFP-hygR mis4-G1487D mip1-R401G* | 2E-G |
| 2 | *h-* | 2H |
| 2729 | *h- mis4-G1487D* | 2H |
| 12167 | *h- mis4-G1487D gtr1-S16N-kanR* | 2H |
| 12179 | *h- mis4-G1487D gtr1-Q61L-kanR* | 2H |
| 12363 | *h- cdc11-GFP-kanR ndc80-GFP-kanR mis6-RFP-hygR mis4-G1487D* | 2-sup1 |
| 12365 | *h- cdc11-GFP-kanR ndc80-GFP-kanR mis6-RFP-hygR* | 2-sup2 |
| 12363 | *h- cdc11-GFP-kanR ndc80-GFP-kanR mis6-RFP-hygR mis4-G1487D* | 2-sup2 |
| 12393 | *h- cdc11-GFP-kanR ndc80-GFP-kanR mis6-RFP-hygR mis4-G1487D mip1-R401G* | 2-sup2 |
| 12634 | *h90 ade6*Δ::*LacO his7+-LacI* | 2-sup3A |
| 12635 | *h90 ade6*Δ::*LacO his7+-LacI mis4-G1487D* | 2-sup3A |
| 12648 | *h90 ade6*Δ::*LacO his7+-LacI mip1-R401G* | 2-sup3A |
| 12653 | *h90 ade6*Δ::*LacO his7+-LacI mis4-G1487D mip1-R401G* | 2-sup3A |
| 2 | *h-* | 2-sup3B |
| 2729 | *h- mis4-G1487D* | 2-sup3B |
| 10874 | *h- mis4-G1487D mip1-R401G* | 2-sup3B |
| 11002 | *h- mip1-R401G* | 2-sup3B |
| 2 | *h-* | 2-sup3C |
| 2729 | *h- mis4-G1487D* | 2-sup3C |
| 2729 | *h-mis4-G1487D* | 2-sup4 |
| 3985 | *h- cdc10-129* | 3B |
| 6392 | *h+ cdc10-129 rad21-FLAG3-kanR* | 3B |
| 8148 | *h+ cdc10-129 rad21-FLAG3-kanR mis4-G1487D* | 3B |
| 11287 | *h+ cdc10-129 rad21-FLAG3-KanR mis4-G1487D mip1-R401G* | 3B |
| 3985 | *h- cdc10-129* | 3C |
| 9748 | *h- cdc10-129 leu1-32 mis4-EGFP-LEU2* | 3C |
| 11357 | *h- cdc10-129 leu1-32 mis4-EGFP-LEU2 mip1-R401G* | 3C |
| 3985 | *h- cdc10-129* | 3D |
| 6392 | *h+ cdc10-129 rad21-FLAG3-KanR* | 3D |
| 11283 | *h+ cdc10-129 rad21-FLAG3-KanR mip1-R401G* | 3D |
| 11326 | *h- psm3-K105NK106N pph3*Δ::*hygR* | 3-sup1A |
| 11395 | *h- psm3-K105NK106N pph3*Δ::*hygR mip1-R401G* | 3-sup1A |
| 2 | *h-* | 3-sup1B |
| 2729 | *h- mis4-G1487D* | 3-sup1B |
| 3985 | *h- cdc10-129* | 4A |
| 9748 | *h- cdc10-129 leu1-32 mis4-EGFP-LEU2* | 4A |
| 3984 | *h- cdc10-129 rad21-9PK-kanR* | 4A |
| 3985 | *h- cdc10-129* | 4B |
| 3984 | *h- cdc10-129 rad21-9PK-kanR* | 4B |
| 10976 | *h- cdc10-129 FLAG-tor2-kanR* | 4B |
| 10984 | *h- cdc10-129 FLAG-tor2-kanR rad21-9PK-kanR* | 4B |
| 2 | *h-* | 4C |
| 10918 | *h- FLAG-tor2-kanR mip1-GFP-hygR* | 4C |
| 10940 | *h- FLAG-tor2-kanR* | 4C |
| 3985 | *h- cdc10-129* | 4-sup1A |
| 9748 | *h- cdc10-129 leu1-32 mis4-EGFP-LEU2* | 4-sup1A |
| 11357 | *h- cdc10-129 leu1-32 mis4-EGFP-LEU2 mip1-R401G* | 4-sup1A |
| 3984 | *h- cdc10-129 rad21-9PK-kanR* | 4-sup1A |
| 3985 | *h- cdc10-129* | 4-sup1A |
| 11029 | *h- cdc10-129 rad21-9PK-kanR mip1-R401G* | 4-sup1A |
| 3984 | *h- cdc10-129 rad21-9PK-kanR* | 4-sup1BC |
| 10976 | *h- cdc10-129 FLAG-tor2-kanR* | 4-sup1BC |
| 10984 | *h- cdc10-129 FLAG-tor2-kanR rad21-9PK-kanR* | 4-sup1BC |
| 11370 | *h- cdc10-129 FLAG-tor2-kanR rad21-9PK-kanR mip1-R401G* | 4-sup1BC |
| 3985 | *h- cdc10-129* | 5A-B |
| 3984 | *h- cdc10-129 rad21-9PK-kanR* | 5A-B |
| 11029 | *h- cdc10-129 rad21-9PK-kanR mip1-R401G* | 5A-B |
| 3985 | *h- cdc10-129* | 5A, 5C |
| 9748 | *h- cdc10-129 leu1-32 mis4-EGFP-LEU2* | 5A, 5C |
| 11357 | *h- cdc10-129 leu1-32 mis4-EGFP-LEU2 mip1-R401G* | 5A, 5C |
| 12287 | *h- mis4-GFP-hygR* | 5-sup1A |
| 12287 | *h- mis4-GFP-hygR* | 5-sup1B |
| 12289 | *h- mis4-GFP-hygR mip1-R401G* | 5-sup1B |
| 12220 | *h- mis4-GFP-hygR mip1-Y533A-13myc* | 5-sup1B |
| 12222 | *h- mis4-GFP-hygR mip1-13myc* | 5-sup1B |
| 3985 | *h- cdc10-129* | 5-sup1C |
| 6393 | *h-cdc10-129 rad21FLAG3-kanR* | 5-sup1C |
| 11284 | *h- cdc10-129 rad21FLAG3-kanR mip1-R401G* | 5-sup1C |
| 11726 | *h- cdc10-129 rad21FLAG3-kanR mip1-Y533A-13myc-kanR* | 5-sup1C |
| 11728 | *h- cdc10-129 rad21FLAG3-kanR mip1-13myc-kanR* | 5-sup1C |
| 3985 | *h- cdc10-129* | 5-sup2AB |
| 9748 | *h- cdc10-129 leu1-32 mis4-EGFP-LEU2* | 5-sup2AB |
| 10289 | *h- cdc10-129 leu1-32 mis4-EGFP-LEU2 pef1*Δ::*hygR* | 5-sup2AB |
| 3985 | *h- cdc10-129* | 5-sup2C |
| 3984 | *h- cdc10-129 rad21-9PK-kanR* | 5-sup2C |
| 6228 | *h- cdc10-129 ura4-D18 pef1*Δ::*ura4+ rad21-9PK-kanR* | 5-sup2C |
| 3985 | *h- cdc10-129* | 5-sup2D |
| 6393 | *h-cdc10-129 rad21FLAG3-kanR* | 5-sup2D |
| 11284 | *h- cdc10-129 rad21FLAG3-kanR mip1-R401G* | 5-sup2D |
| 6360 | *h- cdc10-129 ura4-D18 rad21FLAG3-kanR pef1*Δ::*ura4+* | 5-sup2D |
| 2 | *h-* | 5-sup2E |
| 11478 | *h- psk1-13myc-hygR* | 5-sup2E |
| 11744 | *h+ pef1*Δ::*natR psk1-13myc-hygR* | 5-sup2E |
| 2 | *h-* | 5-sup2F |
| 10369 | *h- pef1*Δ::*natR* | 5-sup2F |
| 11002 | *h- mip1-R401G* | 5-sup2F |
| 11456 | *h- pef1*Δ::*natR mip1-R401G* | 5-sup2F |
| 2 | *h-* | 5-sup3A |
| 2729 | *h-mis4-G1487D* | 5-sup3A |
| 11415 | *h- mis4-G1487D psk1*Δ*::hygR* | 5-sup3A |
| 12240 | *h- mis4-G1487D sck1*Δ*::kanR* | 5-sup3A |
| 12234 | *h- mis4-G1487D sck2*Δ*::kanR* | 5-sup3A |
| 10874 | *h- mis4-G1487D mip1-R401G* | 5-sup3A |
| 12269 | *h- mis4-G1487D sck1*Δ*::kanR sck2*Δ*::kanR* | 5-sup3A |
| 12298 | *h- mis4-G1487D psk1*Δ*::hygR sck1*Δ*::kanR* | 5-sup3A |
| 12267 | *h- mis4-G1487D psk1*Δ*::hygR sck2*Δ*::kanR* | 5-sup3A |
| 12296 | *h- mis4-G1487D psk1*Δ*::hygR sck1*Δ*::kanR sck2*Δ*::kanR* | 5-sup3A |
| 3985 | *h- cdc10-129* | 5-sup3B |
| 6392 | *h+ cdc10-129 rad21-FLAG3-kanR* | 5-sup3B |
| 11283 | *h+ cdc10-129 rad21-FLAG3-KanR mip1-R401G* | 5-sup3B |
| 12284 | *h+ cdc10-129 rad21-FLAG3-kanR sck2*Δ*::kanR* | 5-sup3B |
| 12290 | *h+ cdc10-129 rad21-FLAG3-kanR sck1*Δ*::kanR* | 5-sup3B |
| 12307 | *h+ cdc10-129 rad21-FLAG3-kanR sck1*Δ*::kanR sck2*Δ*::kanR* | 5-sup3B |
| 2 | *h-* | 5-sup3C |
| 2729 | *h-mis4-G1487D* | 5-sup3C |
| 12240 | *h- mis4-G1487D sck1*Δ*::kanR* | 5-sup3C |
| 12234 | *h- mis4-G1487D sck2*Δ*::kanR* | 5-sup3C |
| 12269 | *h- mis4-G1487D sck1*Δ*::kanR sck2*Δ*::kanR* | 5-sup3C |
| 3985 | *h- cdc10-129* | 5-sup3D |
| 6392 | *h+ cdc10-129 rad21-FLAG3-kanR* | 5-sup3D |
| 11283 | *h+ cdc10-129 rad21-FLAG3-KanR mip1-R401G* | 5-sup3D |
| 12284 | *h+ cdc10-129 rad21-FLAG3-kanR sck2*Δ*::kanR* | 5-sup3D |
| 12290 | *h+ cdc10-129 rad21-FLAG3-kanR sck1*Δ*::kanR* | 5-sup3D |
| 12307 | *h+ cdc10-129 rad21-FLAG3-kanR sck1*Δ*::kanR sck2*Δ*::kanR* | 5-sup3D |
| 9748 | *h- cdc10-129 leu1-32 mis4-EGFP-LEU2* | 5-sup3E |
| 12302 | *h- cdc10-129 leu1-32 mis4-EGFP-LEU2 sck1*Δ*::kanR* | 5-sup3E |
| 12306 | *h- cdc10-129 leu1-32 mis4-EGFP-LEU2 sck2*Δ*::kanR* | 5-sup3E |
| 12309 | *h- cdc10-129 leu1-32 mis4-EGFP-LEU2 sck1*Δ*::kanR sck2*Δ*::kanR* | 5-sup3E |
| 2 | *h-* | 6A |
| 2729 | *h- mis4-G1487D* | 6A |
| 11164 | *h- mis4-G1487D psm1-S1022A* | 6A |
| 11897 | *h- mis4-G1487D-S183A* | 6A |
| 11985 | *h- mis4-G1487D-S183A psm1-S1022A* | 6A |
| 2 | *h-* | 6A |
| 2729 | *h- mis4-G1487D* | 6A |
| 11128 | *h- mis4-G1487D psm1-S1022D* | 6A |
| 11857 | *h- mis4-G1487D-S183E* | 6A |
| 11987 | *h- mis4-G1487D-S183E psm1-S1022D* | 6A |
| 2729 | *h- mis4-G1487D* | 6B |
| 11985 | *h- mis4-G1487D-S183A psm1-S1022A* | 6B |
| 11987 | *h- mis4-G1487D-S183E psm1-S1022D* | 6B |
| 12363 | *h- cdc11-GFP-kanR ndc80-GFP-kanR mis6-RFP-hygR mis4-G1487D* | 6C |
| 12365 | *h- cdc11-GFP-kanR ndc80-GFP-kanR mis6-RFP-hygR* | 6C |
| 12626 | *h- cdc11-GFP-kanR ndc80-GFP-kanR mis6-RFP-hygR mis4-G1487D-S183A psm1-S1022A* | 6C |
| 12644 | *h- cdc11-GFP-kanR ndc80-GFP-kanR mis6-RFP-hygR mis4-G1487D-S183E psm1-S1022D* | 6C |
| 12626 | *h- cdc11-GFP-kanR ndc80-GFP-kanR mis6-RFP-hygR mis4-G1487D-S183A psm1-S1022A* | 6D |
| 12644 | *h- cdc11-GFP-kanR ndc80-GFP-kanR mis6-RFP-hygR mis4-G1487D-S183E psm1-S1022D* | 6D |
| 2 | *h-* | 6E-6sup1 |
| 2729 | *h- mis4-G1487D* | 6E-6sup1 |
| 11985 | *h- mis4-G1487D-S183A psm1-S1022A* | 6E-6sup1 |
| 11987 | *h- mis4-G1487D-S183E psm1-S1022D* | 6E-6sup1 |
| 2 | *h-* | 6-sup2A |
| 11326 | *h- pph3::hygR psm3-K105NK106N* | 6-sup2A |
| 12621 | *h- pph3::hygR psm3-K105NK106N mis4-S183A psm1-S1022A* | 6-sup2A |
| 12617 | *h- pph3::hygR mis4-S183A psm1-S1022A* | 6-sup2A |
| 12613 | *h- psm3-K105NK106N mis4-S183A psm1-S1022A* | 6-sup2A |
| 6095 | *h+ pph3::hygR* | 6-sup2A |
| 4573 | *h- psm3-K105NK106N* | 6-sup2A |
| 11326 | *h- pph3::hygR psm3-K105NK106N* | 6-sup2BC |
| 11395 | *h- pph3::kanR psm3K105NK106N mip1-R401G* | 6-sup2BC |
| 11519 | *h- pph3::hygR psm3-K105NK106N psm1-S1022A* | 6-sup2BC |
| 11918 | *h- pph3::kanR psm3K105NK106N mis4-S183A* | 6-sup2BC |
| 12621 | *h- pph3::hygR psm3-K105NK106N mis4-S183A psm1-S1022A* | 6-sup2BC |
| 12837 | *h-/h- ade6-210/ade6-216* | 7 |
| 12841 | *h-/h- ade6-210/ade6-216* | 7 |
| 12727 | *h+ ade6-704 CM3112* | 7 |
| 12765 | *h+ ade6-704 CM3112 mip1-R401G* | 7 |
| 12822 | *h+ ade6-704 CM3112 psm1-S1022A* | 7 |
| 12845 | *h+ ade6-704 CM3112 psm1-S1022D* | 7 |
| 12826 | *h+ ade6-704 CM3112 psm1-S1022D mis4-S183E* | 7-sup1 |
| 12831 | *h+ ade6-704 CM3112 mis4-S183A* | 7-sup1 |
| 12833 | *h+ ade6-704 CM3112 mis4-S183E* | 7-7sup1 |
| 12863 | *h+ ade6-704 CM3112 psm1-S1022A mis4-S183A* | 7-sup1 |
| 11589 | *mat1-M-smt0* | 8 |
| 11654 | *mat1-M-smt0 mis4-G1487D* | 8 |
| 11628 | *mat1-M-smt0 mip1-R401G* | 8 |
| 11626 | *mat1-M-smt0 mis4-G1487D mip1-R401G* | 8 |
